# BioASQ-QA: A manually curated corpus for Biomedical Question Answering

**DOI:** 10.1101/2022.12.14.520213

**Authors:** Anastasia Krithara, Anastasios Nentidis, Konstantinos Bougiatiotis, Georgios Paliouras

## Abstract

The BioASQ question answering (QA) benchmark dataset contains questions in English, along with golden standard (reference) answers and related material. The dataset has been designed to reflect real information needs of biomedical experts and is therefore more realistic and challenging than most existing datasets. Furthermore, unlike most previous QA benchmarks that contain only exact answers, the BioASQ-QA dataset also includes ideal answers (in effect summaries), which are particularly useful for research on multi-document summarization. The dataset combines structured and unstructured data. The material linked with each question comprise documents and snippets, which are useful for Information Retrieval and Passage Retrieval experiments, as well as concepts that are useful in concept-to-text Natural Language Generation. Researchers working on paraphrasing and textual entailment can also measure the degree to which their methods improve the performance of biomedical QA systems. Last but not least, the dataset is continuously extended, as the BioASQ challenge is running and new data are generated.

## Background & Summary

More than 2 articles are published in biomedical journals every minute, leading to MEDLINE^1^ currently comprising more than 32 million articles, while the number and size of non-textual biomedical data sources also increases rapidly. As an example, since the outbreak of the COVID-19 pandemic, there has been an explosion of new scientific literature about the disease and the virus that causes it, with about 10,000 new COVID-19 related articles added each month^1^. This wealth of new knowledge plays a central role in the progress achieved in biomedicine and its impact on public health, but it is also overwhelming for the biomedical expert. Ensuring that this knowledge is used for the benefit of the patients in a timely manner is a demanding task.

BioASQ^2^ (Biomedical Semantic Indexing and Question Answering) pushes research towards highly precise biomedical information access systems through a series of evaluation campaigns, in which systems from teams around the world compete. BioASQ campaigns run annually since 2012, providing data, open-source software and a stable evaluation environment for the participating systems. In the last ten years that the challenge has been running, around 100 different universities and companies, from all continents, have participated in BioASQ, providing a competitive, but also synergetic ecosystem. The fact that the participants of the BioASQ challenges are all working on the same benchmark data, facilitates significantly the exchange and fusion of ideas and eventually accelerates progress in the field. The ultimate goal is to lead biomedical information access systems to the maturity and reliability required by biomedical researchers.

BioASQ comprises two main tasks. In Task A systems are asked to automatically assign MeSH^3^ terms to biomedical articles, thus assisting the indexing of biomedical literature. Task B focuses on obtaining precise and comprehensible answers to biomedical research questions. The systems that participate in Task B are given English questions written by biomedical experts that reflect real-life information needs. For each question, the systems are required to return relevant articles, snippets of the articles, concepts from designated ontologies, RDF triples from Linked Life Data^4^, an ‘exact’ answer (e.g., a disease or symptom), and a paragraph-sized summary answer. Hence, this task combines traditional information retrieval, with question answering from text and structured data, as well as multi-document text summarization.

One of the main tangible outcomes of BioASQ is its benchmark datasets. The BioASQ-QA dataset that is generated for Task B, contains questions in English, along with golden standard (reference) answers and supporting material. The BioASQ data is more realistic and challenging than most existing datasets for question answering. In order to achieve this, BioASQ employs a team of trained experts, who provide annually a set of around 500 questions from their specialized field of expertise. Figure 1 provides the lifecycle of the BioASQ dataset creation, which is presented in detail in the following sections. Using this process, a set of 4721 questions and answers have been generated so far, constituting a unique resource for the development of QA systems.

**Figure 1.**
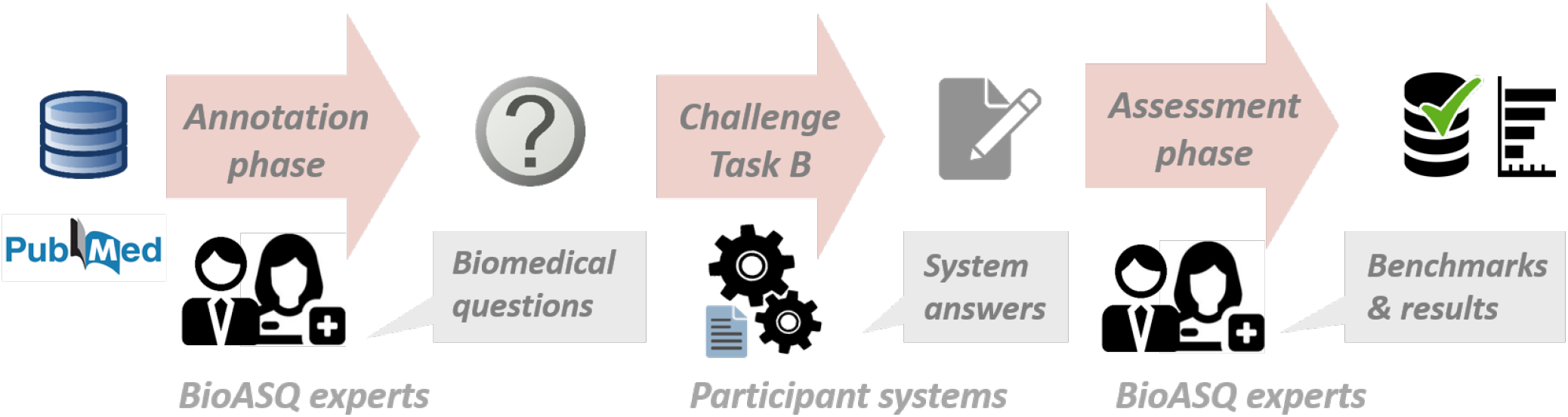
During the annotation phase of the BioASQ, the experts compose biomedical questions. The participating systems provide answers in the challenge. Finally, in the assessment phase, the experts manually assess the system responses and refine and extend the dataset.

## Methods

### The BioASQ infrastructure and ecosystem

Figure 2 summarises the main components of the BioASQ infrastructure, as well as key stakeholders in the related ecosystem. The BioASQ infrastructure includes tools for annotating data, tools for assessing the results of participating systems, benchmark repositories, evaluation services, etc. The infrastructure allows challenge participants to access training and test data, submit their results and be informed about the performance of their systems, in comparison to other systems. The BioASQ infrastructure is also used by the experts during the creation of the benchmark datasets and helps improve the quality of the data. In the following subsections, the different components of the BioASQ ecosystems are described.

**Figure 2.**
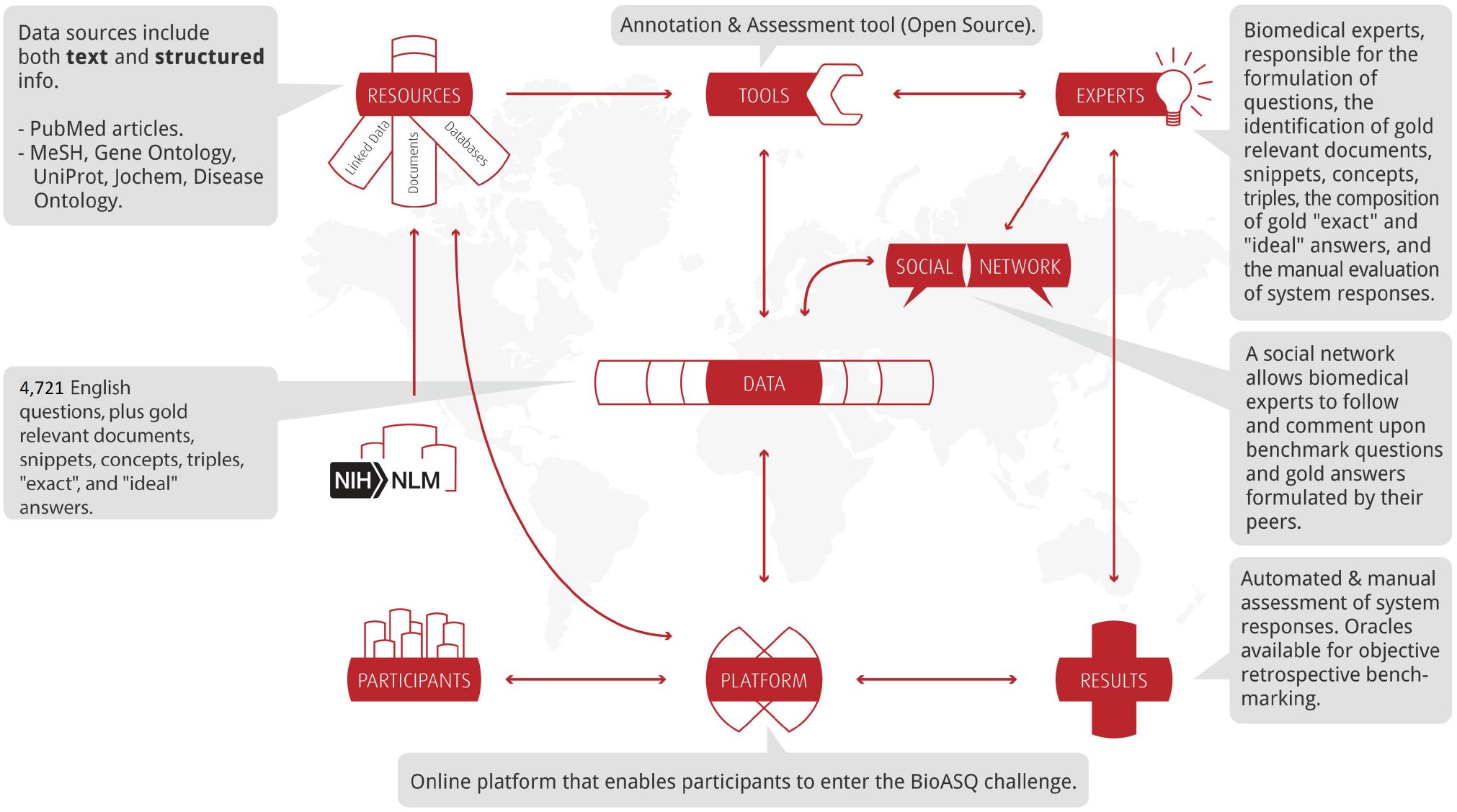
The BioASQ infrastructure and ecosystem

### Expert team

As the goal of BioASQ is to reflect real information needs of biomedical experts, their involvement was necessary in the creation of the dataset. The biomedical expert team of BioASQ was first established in 2012, but has changed through the years. Several experts were considered at that time, from a variety of institutions across Europe. The final selection of the experts was based on the need to cover the broad biomedical scientific field, representing as much as possible, medicine, biosciences and bioinformatics. The members of the biomedical team hold positions in universities, hospitals or research institutes in Europe. Their primary research interests include: cardiovascular endocrinology, psychiatry, psychophysiology, pharmacology, drug repositioning, cardiac remodeling, cardiovascular pharmacology, computational genomics, pharmacogenomics, comparative genomics, molecular evolution, proteomics, mass spectometry, protein evolution, clinical information retrieval from electronic health records, and clinical practice guidelines. In total 21 experts have contributed to the creation of the dataset, 7 of whom have been involved most actively. The main job of the biomedical expert team is the creation of the QA benchmark dataset, using an annotation tool provided by BiOASQ. With the use of the tool, the experts can set their questions and retrieve relevant documents and snippets from MEDLINE. Additionally, the biomedical expert team assesses the responses of the participating systems. In addition to scoring the systems’ answers, during this process the experts have the opportunity to enrich and modify the gold material that they have provided, thus improving the quality of the benchmark dataset.

Regular physical and virtual meetings are organised with the experts. Partly, these meetings aim to train the new members of the team and inform the existing ones about changes that have happened. In particular, the goals of the training sessions are as follows:

- Familiarization with the annotation and assessment tools used during the formulation and assessment of biomedical questions respectively. This step also involves familiarization of the experts with the specific types of questions used in the challenge, i.e. factoid, yes/no, list and summary questions. At the same time, the experts provide feedback and help shaping of the BioASQ tools themselves.
- Familiarization with the resources used in BioASQ, both MEDLINE and various structured sources. The aim is to help the experts understand the data provided by theses sources, in response to different questions they may formulate.
- Resolution of issues that come up during the question composition and assessment tasks. This is a continuous process that extends beyond the training sessions. Continuous support is provided to the experts, while the experts can also interact with each other and provide feedback on the data being created.

### Data selection

The QA benchmark is based primarily on documents indexed by *MEDLINE*. In addition a wide range of biomedical concepts are drawn from ontologies and linked data that describe different facets of the domain. The selected resources follow commonly used *drug-target-disease* triangle, which defines the prime information axes for medical investigations. The main principle is shown in Figure 3.

**Figure 3.**
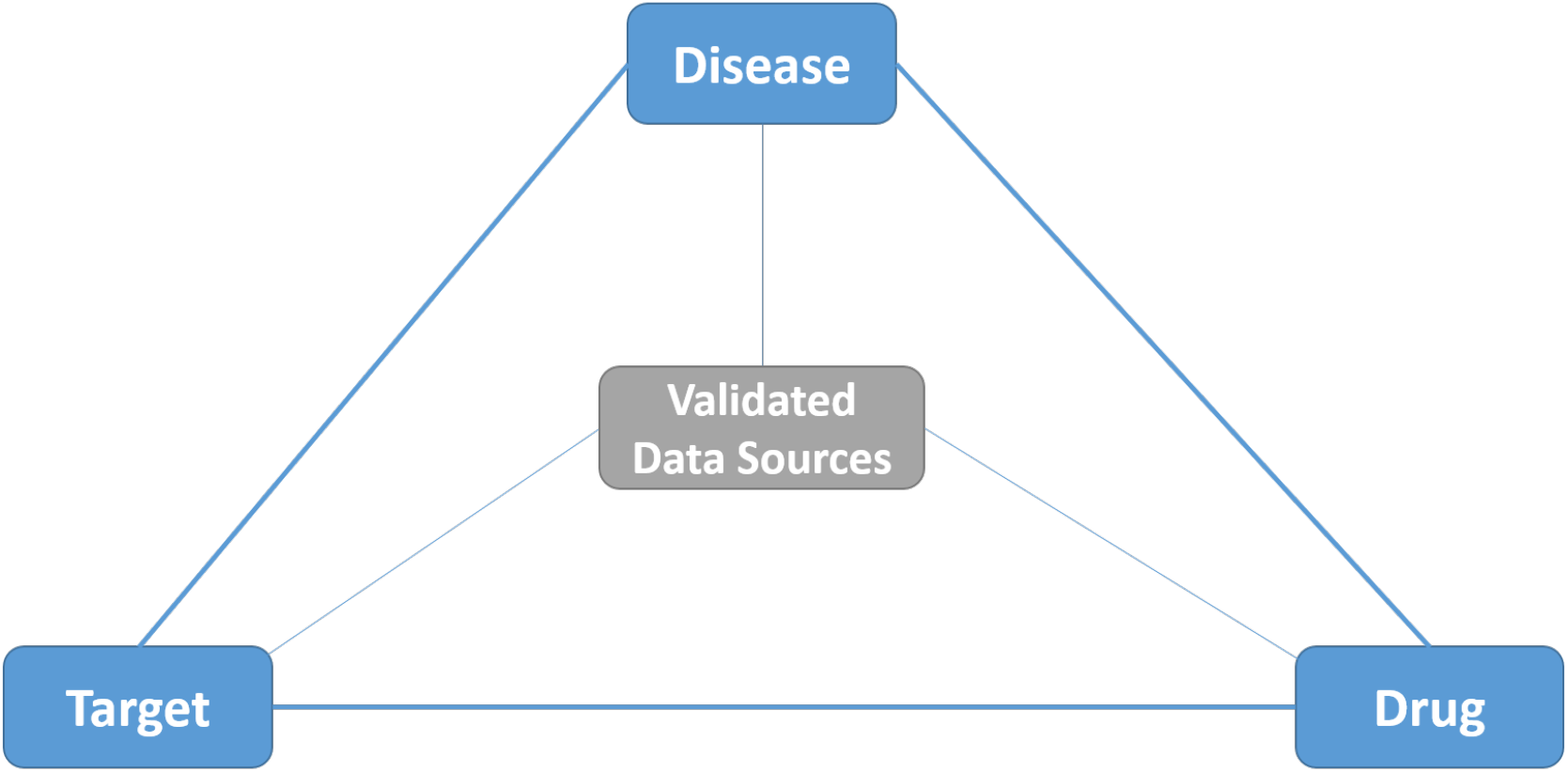
The drug-target-disease triangle, adopted in *BioASQ*.

This *“knowledge-triangle*” supports the conceptual linking of biomedical knowledge databases and related resources. Based on this, systems can address questions, linking natural language questions with relevant ontology concepts. In this context, the following resources have been selected for BioASQ.

#### Drugs

**Jochem**^2^, the Joint Chemical Dictionary, is a dictionary for the identification of small molecules and drugs in text, combining information from UMLS, MeSH, ChEBI, DrugBank, KEGG, HMDB, and ChemIDplus. Given the variety and the population of the different resources in it, Jochem is currently one of the largest biomedical resources for drugs and chemicals.

#### Targets

**Gene Ontology** (GO)^3,4^ is currently the most successful case of ontology use in bioinformatics and provides a controlled vocabulary to describe functional aspects of gene products. The ontology covers three domains: cellular component, molecular function, and biological process.

#### Universal Protein Resource

(UniProt^5^) provides a comprehensive, high-quality and freely accessible resource of protein sequence and functional information. Its protein knowledge base consists of two sections: SwissProt, which is manually annotated and reviewed, and contains more than 500 thousand sequences, and TrEMBL, which is automatically annotated and is not reviewed, and contains a few million sequences. In BioASQ the SwissProt component of UniProt is used.

#### Diseases

**Disease Ontology** (DO)^5^ contains data associating genes with human diseases, using established disease codes and terminologies. Approximately 8,000 inherited, developmental and acquired human diseases are included in the resource. The DO semantically integrates disease and medical vocabularies through extensive cross-mapping and integration of MeSH, ICD, NCI’s thesaurus, SNOMED CT and OMIM disease-specific terms and identifiers.

#### Document Sources

The main source of biomedical literature is NLM’s MEDLINE and is accessible through PubMed and PubMed Central. PubMed, indexes over 34 million citations, while PubMed Central (PMC) provides free access to approximately 8.5 million full-text biomedical and life-science articles.

#### The Medical Subject Headings Hierarchy

(MeSH) is a hierarchy of terms maintained by the US National Library of Medicine (NLM) and its purpose is to provide headings (terms), which can be used to index scientific publications in the life sciences, e.g., journal articles, books, and articles in conference proceedings. The indexed publications may be searched through popular search engines, such as PubMed, using the MeSH headings to filter semantically the results. This retrieval methodology seems to be in some cases beneficial, especially when precision of the retrieved results is important^6^. The primary MeSH terms (called *descriptors*) are organized into 16 trees, and are approximately 30,200. MeSH is the main resource used by PubMed to index the biomedical scientific bibliography in MEDLINE.

#### Linked Data

During the first few years of BioASQ, the Linked Life Data platform was used to identify subject-verb-object triples related to questions. Linked Life Data is a data warehouse that syndicates large volumes of heterogeneous biomedical knowledge in a common data model. It contains more than 10 billion statements. The statements are extracted from 25 biomedical resources, such as PubMed, UMLS, DrugBank, Diseasome, and Gene Ontology. This resource has been abandoned in recent editions of BioASQ, due to issues with the triple selection process.

### Question formulation

The members of the biomedical expert team formulate English questions, reflecting real-life information needs encountered during their work (e.g., in diagnostic research). Figure 4 provides an overview of the most frequent topics covered in the questions generated so far by the experts. Each question is independent of all other questions and is associated with an answer and other supportive information, as explained below.

**Figure 4.**
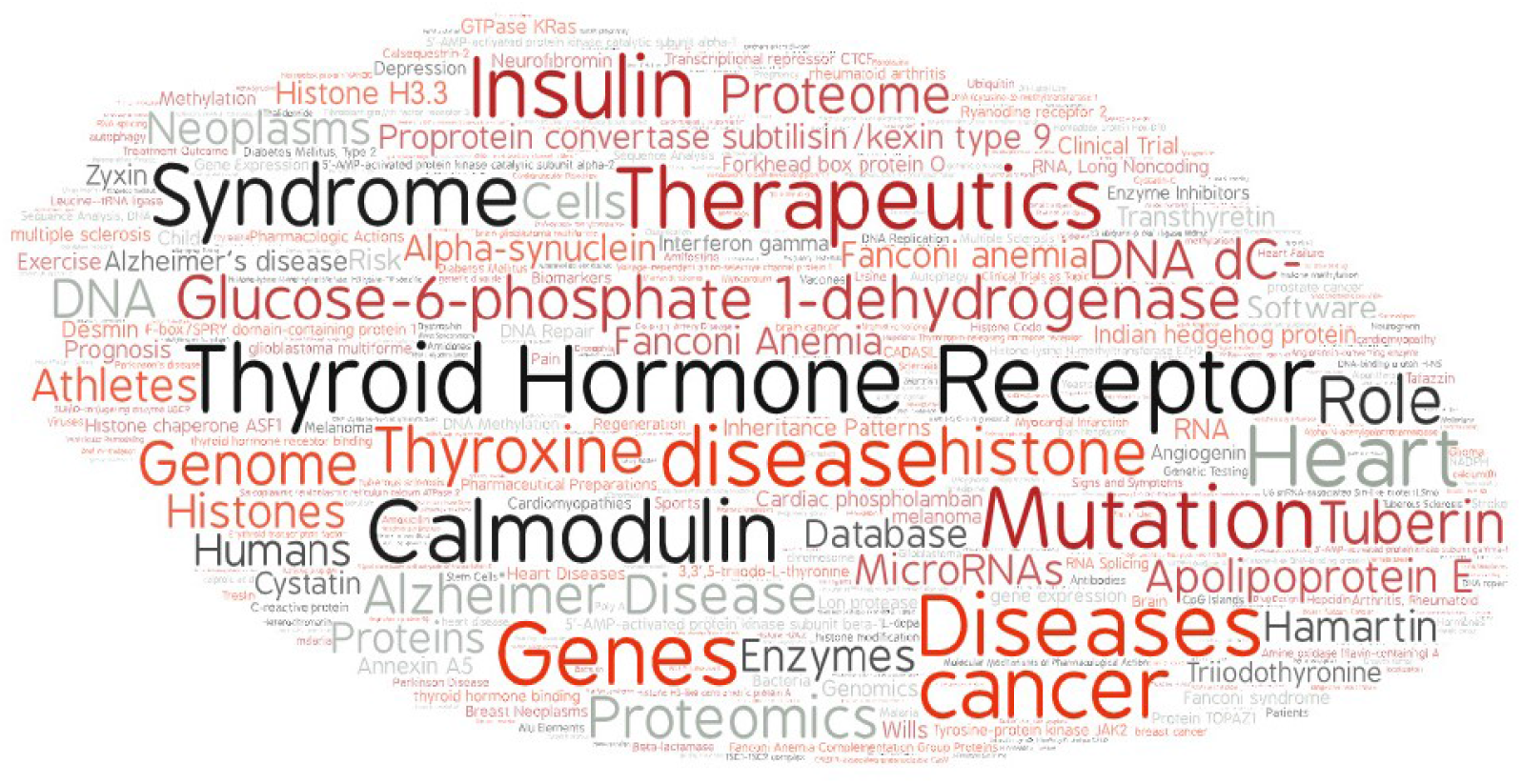
Most frequent topics in the BioASQ questions

In addition to the training sessions mentioned above, guidelines are provided to the BioASQ experts to help them create the questions, reference answers, and other supportive information^7^. The guidelines cover the number and types of questions to be created by the experts, the information sources the experts should consider and how to use them, the types and sizes of the answers, additional supportive information the experts should provide, etc. The experts use the BioASQ annotation tool for this process, which is accessible through a Web interface^6^.

The annotation tool provides the necessary functionality to create questions and select relevant information. The annotation tool is designed to be easy to use, adopting a simple five-step-paradigm: authenticate, search, select, annotate and store. The authentication ensures that each question created by a certain expert can be assigned to this given expert.

The annotation process comprises the following steps:

#### Step 1: Question formulation

The experts formulate an English stand-alone question, reflecting their information needs. Questions may belong to one of the following four categories:

**Yes/no questions:** These are questions that, strictly speaking, require either a “yes” or a “no” as an answer, though of course in practice a longer answer providing additional information is useful. For example, “*Do CpG islands colocalise with transcription start sites?*” is a yes/no question.
**Factoid questions:** These are questions that require a particular entity (e.g., a disease, drug, or gene) as an answer, though again a longer answer is useful. For example, “*Which virus is best known as the cause of infectious mononucleosis?*” is a factoid question.
**List questions:** These are questions that require a *list* of entities (e.g., a list of genes) as an answer; again, in practice additional supportive information is desirable. For example, “*Which are the Raf kinase inhibitors?*” is a list question.
**Summary questions:** These are questions that do not belong in any of the previous categories and can only be answered by producing a short text summarizing the most prominent relevant information. For example, *“How does dabigatran therapy affect aPTT in patients with atrial fibrillation?*” is a summary question. When formulating summary questions, the experts aimed at questions that they can answer in a satisfactory manner with a one-paragraph summary, intended to be read by other experts of the same field.

In all four categories, the experts aim at questions for which a limited number of articles (min. 10, max. 60) are retrieved through PubMed queries. Questions which are controversial or that have no clear answers in the literature are avoided. Moreover, all questions are related to the biomedical domain. For example, in the case of the following two questions:

*Q*_1_: Which are the differences between Hidden Markov Models (HMMs) and Artificial Neural Networks (ANNs)? *Q*_2_: Which are the uses of Hidden Markov Models (HMMs) in gene prediction?

Although HMMs and ANNs are used in the biomedical domain, *Q*_1_ is not suitable for the needs of BioASQ, since there is not a direct indication that it is related to the biomedical domain. On the other hand, *Q*_2_ links to “gene prediction” and is appropriate.

#### Step 2: Relevant terms

A set of terms that are relevant to each question is selected. The set of relevant terms may include terms that are already mentioned in the question, but it may also include synonyms of the question terms, closely related broader and narrower terms etc. For the question “*Do CpG islands colocalise with transcription start sites?*”, the set of relevant terms would most probably include the question terms “*CpG Island*” and “*transcription start site*”, but possibly also other terms, like the synonym “*Transcription Initiation Site*”.

#### Step 3: Information retrieval

Using the selected terms, the BioASQ annotation tool allows the experts to issue queries and retrieve relevant articles through PubMed. More than one query may be associated with each question and each query can be enriched with the advanced search tags of PubMed.^7^. The search window (Figure 5) allows selecting information that is necessary to answer the question. One of the main powers of the annotation tool is that it implements interfaces to different data sources of different types, i.e., unstructured, semi-structured or structured. Given that we cannot expect domain experts to be familiar with Semantic Web standards, such as RDF, the annotation tool also implements an innovative natural language generation method that converts RDF into natural language. The iterative improvement of the annotation tool has led to a framework that is widely accepted by the BioASQ biomedical expert team. Interestingly, a study of the queries used by different experts to answer the same questions made clear that indeed “many roads lead to Rome” „ i.e. different experts will use different queries for the same question.

**Figure 5.**
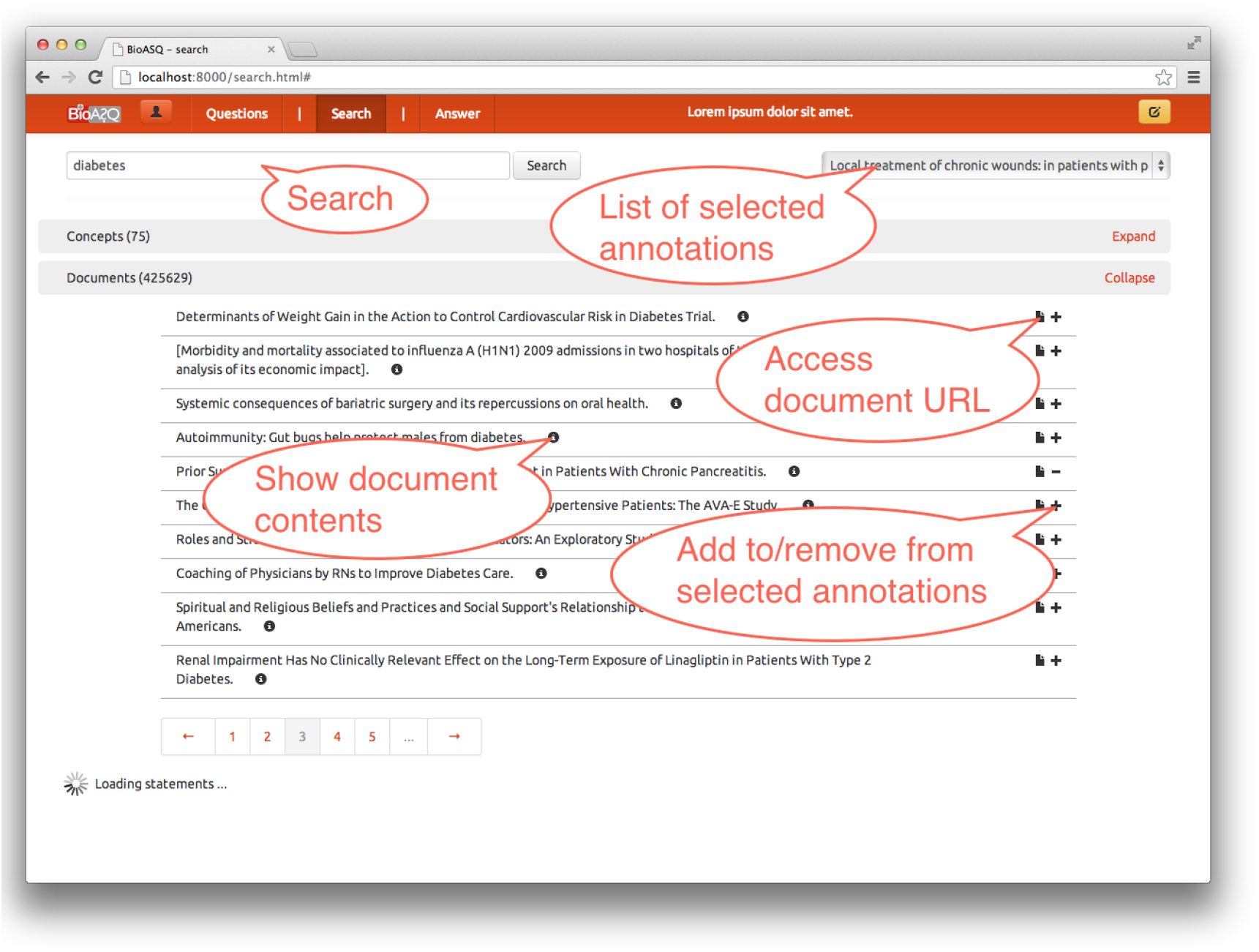
Screenshot of the annotation tool’s search and data selection screen with the section for *document results* expanded.

Returning to the example question “*Do CpG islands colocalise with transcription start sites?*” a query may be “*CpG Island*”*AND “transcription start site*”. Some of the articles retrieved by this query are shown in Table 1.

**Table 1.**
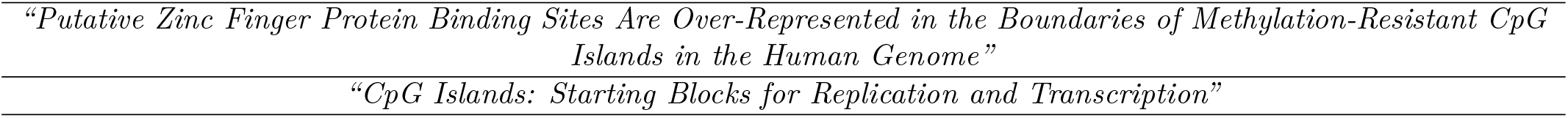
Examples of retrieved articles (only titles shown here, but the annotation tool provides also the abstracts)

#### Step 4: Selection of articles

Based on the results of Step 3, the experts select a set of articles that are sufficient for answering the question. Using the annotation tool, they choose among the retrieved list of articles, the ones that contain relevant information to form an answer.

#### Step 5: Text snippet extraction

Using the articles selected in step 4, the experts mark *every* text snippet (piece of text) out of the articles selected in Step 4. Snippets can be easily extracted using the annotation tool (Figure 6) and may answer the question either fully or partially the question. A text snippet should contain one or more entire and consecutive sentences. If there are multiple snippets that provide the same (or almost the same) information (in the same or in different articles), *all* of them are selected. Examples of relevant snippets are shown in Table 2.

**Figure 6.**
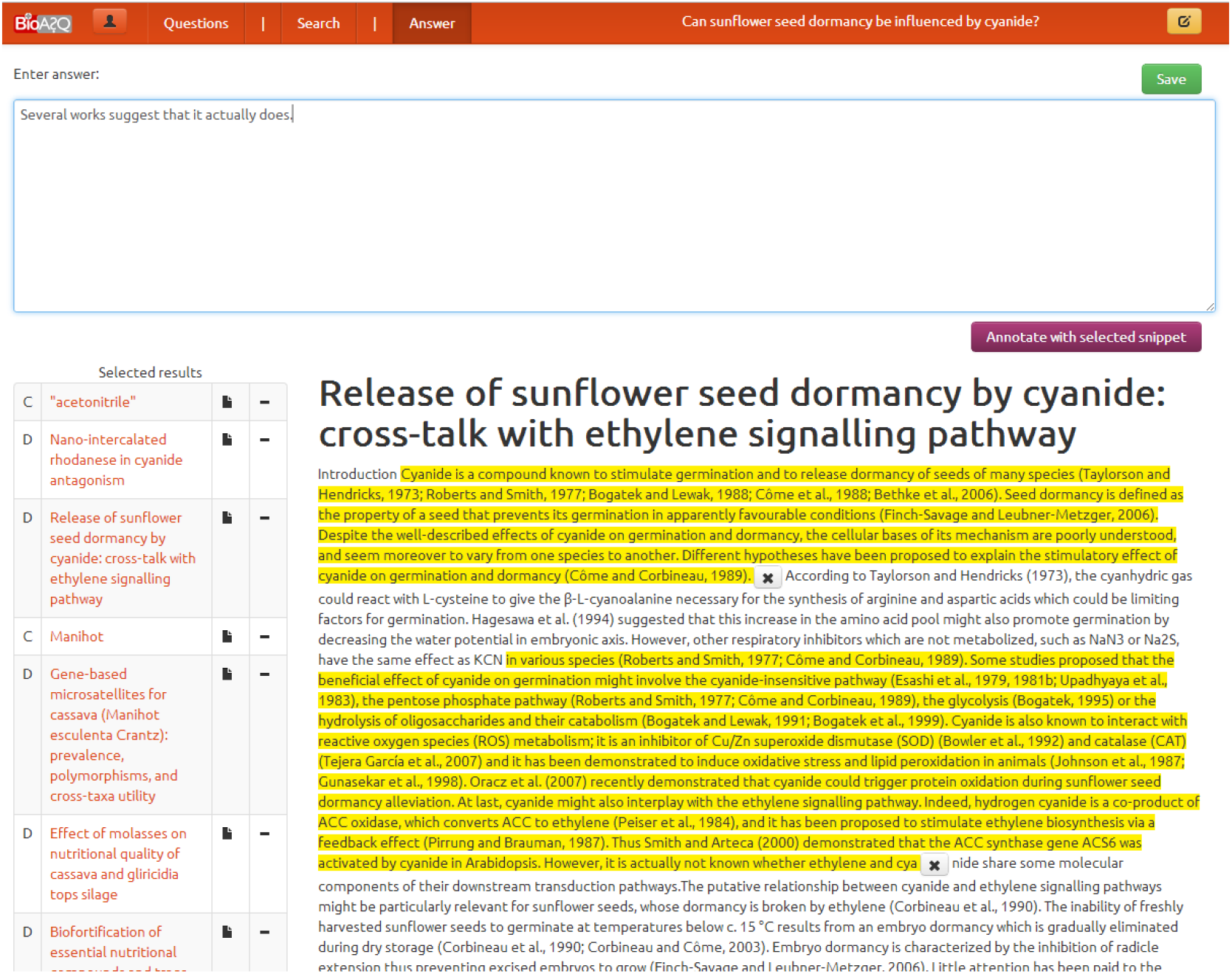
Screenshot of the annotation tool’s snippet annotation process

**Table 2.**
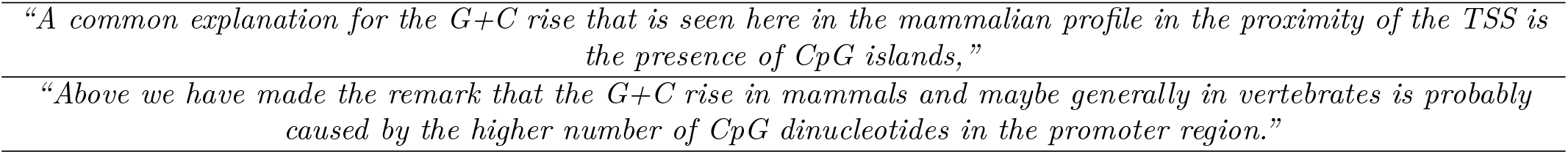
Examples of relevant snippets

#### Step 6: Query revision

If the expert judges that the articles and snippets gathered during steps 2 to 5 are insufficient for answering the question, the process can be repeated. The articles that the expert has already selected can be saved before performing a new search, along with the snippets the expert has already extracted. The query can be revised several times, until the expert feels that the gathered information is sufficient to answer the question. At the end, if the expert judges that the question can still not be answered adequately, the question is discarded.

#### Step 7: Exact answer

In steps 2 to 6, the expert identifies relevant material for answering the question. Given this material, the next step is to formulate the actual answer. For a yes/no question, the exact answer is simply “yes” or “no”. For a factoid question, the exact answer is the name of the entity (e.g., gene, disease) sought by the question; if the entity has several synonyms, the expert provides, to the extent possible, all of its synonyms. For a list question, the exact answer is a list containing the entities sought by the question; if a member of the list has several synonyms, the expert provides again as many of the synonyms as possible. For a summary question, the exact answer is left blank. The exact answers of yes/no, factoid, and list questions should be based on the information of the text snippets that the expert has selected, rather than personal experience.

#### Step 8: Ideal answer

At this final step, the expert formulates what we call an *ideal answer* for the question. The ideal answer should be a one-paragraph text that answers the question in a manner that the expert finds satisfactory. The ideal answer should be written in English, and it should be intended to be read by other experts of the same field. For the example question “*Do CpG islands colocalise with transcription start sites?*”, an ideal answer might be the one shown in Table 3. Again, the ideal answer should be based on the information of the text snippets that the expert has selected, rather than personal experience. The experts, however, are allowed (and should) rephrase or shorten the snippets, order or combine them etc., in order to make the ideal answer more concise and easier to read.

**Table 3.**
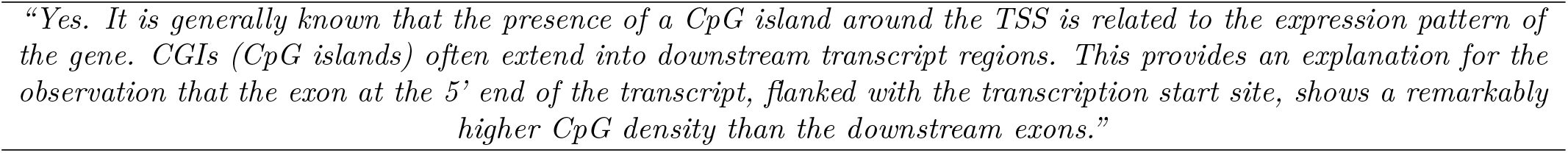
Examples of relevant snippets

Notice that in the example above, the ideal answer provides additional information supporting the exact answer. If the expert feels that the exact answer of a yes/no, factoid, or list question is sufficient and no additional information needs to be reported, the ideal answer can be the same as the exact answer. For summary questions, an ideal answer must always be provided.

Figure 7 presents the distribution of questions created each year of the challenge. During the years, there is an increase in the number of factoid questions, and a decrease in the number of list questions. The possible reason is that it is more difficult to find the material (i.e. articles and snippets) that are sufficient for answering a factoid question than a list question. Table 4 presents the different versions of the BioASQ-QA dataset, including the number of questions, and the average number of documents and snippets^8^. Each version of the training dataset enriches its previous version with the new questions created the respective year.

**Figure 7.**
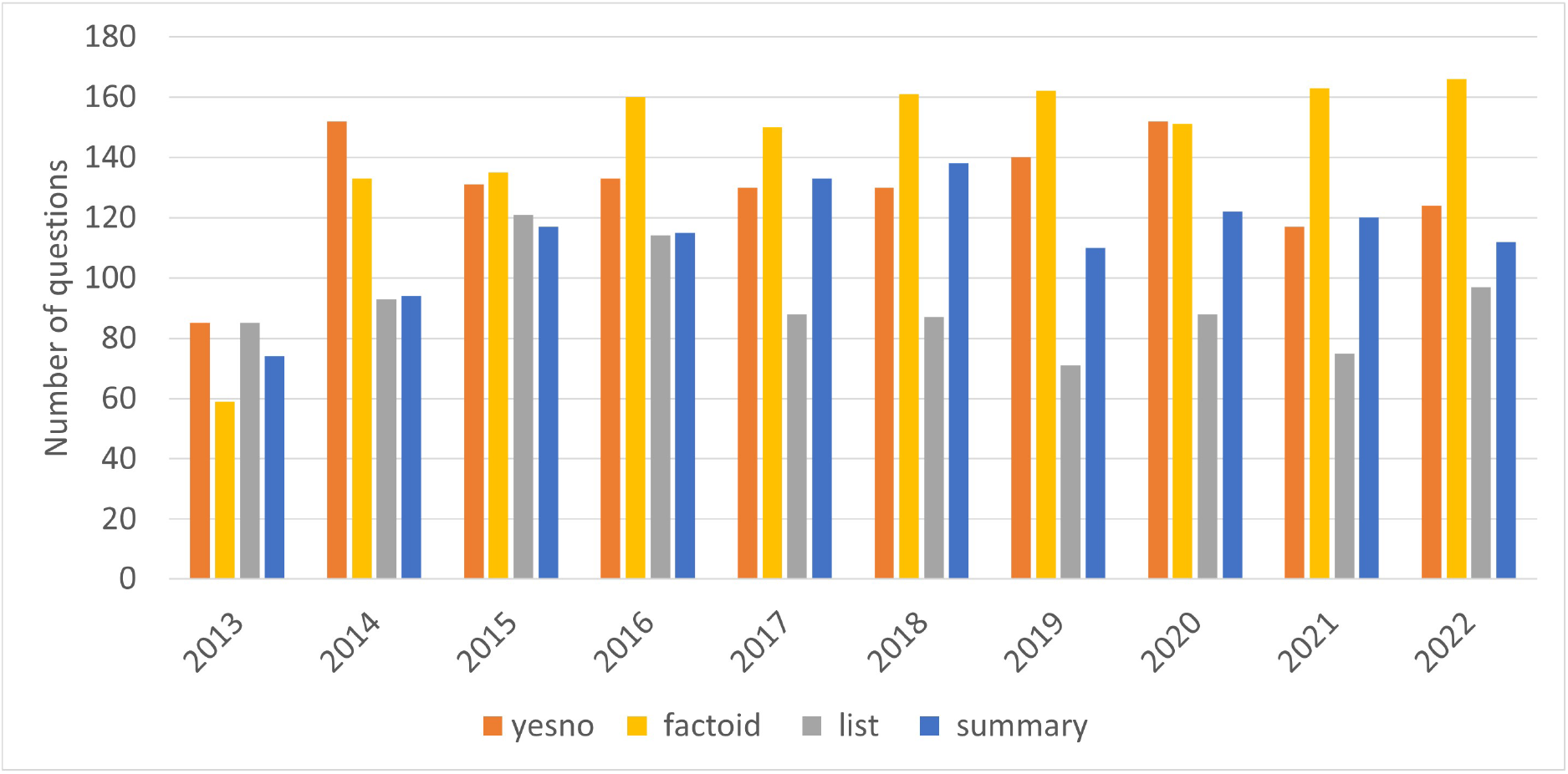
Distribution of types of questions per year

**Table 4.**
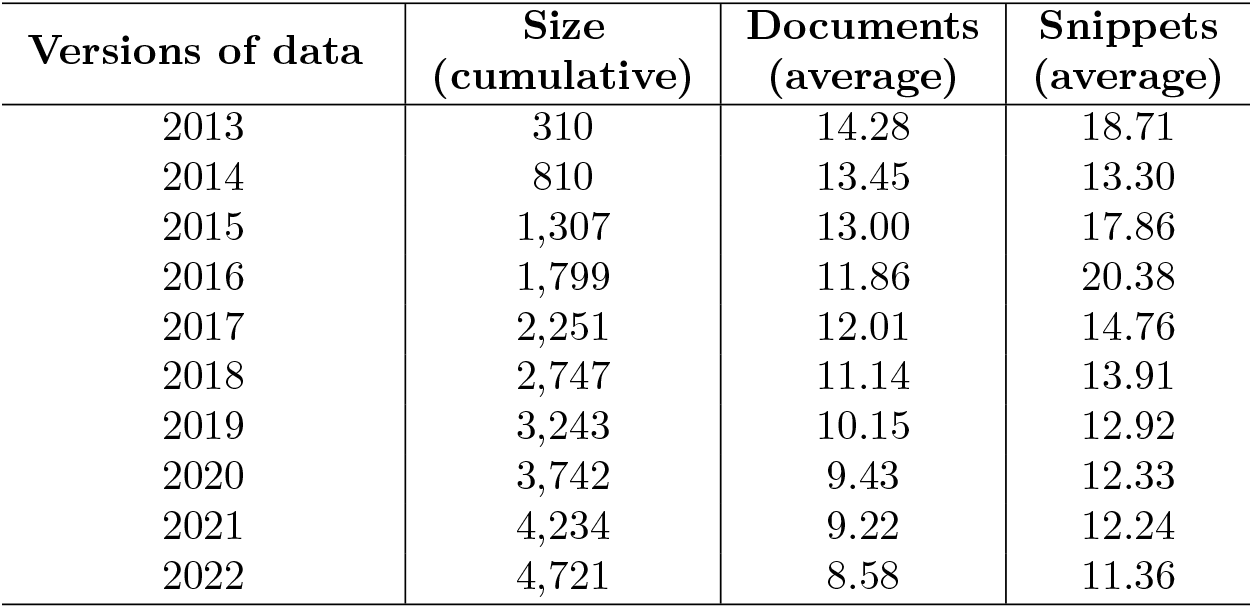
The different versions of the BioASQ-QA dataset, as it has evolved over the years of the challenge.

During the years that BioASQ has been running, three significant changes have taken place, in response to feedback obtained by the experts and the challenge participants:

- Since BioASQ 3 (2015), the focus of the experts is only on relevant articles and their contents. In other words, the experts do not provide relevant concepts or statements, as it was found cumbersome and led to questionable results. Nevertheless, concepts are included in the gold dataset, as they are added by the systems and assessed by the experts in the assessment phase).
- Since BioSQ 4 (2016) only a sufficient set of articles, that allow the answer to be found with confidence, is requested by the experts. This is again in contrast to earlier years, where the experts were asked to identify all relevant articles; something that proved to be unrealistic. Again, if the participating systems retrieve more relevant documents, not identified in the annotation phase by the experts, these are added in the gold dataset, during the assessment phase.
- In early versions of the challenge, we considered using full-text articles from PubMed Central (PMC). Given the small percentage of the overall literature that appears in PMC, since BioASQ 4 (2016) we decided to restrict the challenge to article abstracts only.

### Assessment

Following each round of the challenge, the answers of the participating systems are collected and assessed. Exact answers can be assessed automatically against the golden answers provided by the experts during the annotation phase. However, the ‘ideal’ answers are assessed manually by the experts. In fact each expert gets to assess the answers to the questions they have created, in terms of *information recall* (does the ‘ideal’ answer reports all the necessary information?), *information precision* (does the answer contain only relevant information?), *information repetition* (does the ‘ideal’ answer avoid repeating the same information multiple times? e.g., when sentences of the ‘ideal’ answer that have been extracted from different articles convey the same information), and *readability* (is the ‘ideal’ answer easily readable and fluent?). A 1 to 5 scale is used in all four criteria (1 for ‘very poor’, 5 for ‘excellent’).

The assessment tool is designed to be a companion to the annotation tool and is implemented by reusing most of its functionality. The tool can also be used to perform an inter-annotator agreement study. In that case, domain experts are provided with answers generated by other (anonymous) domain experts and are asked to evaluate them.

The design of the interface is such that the users can always see the answers/annotations only to questions that they are asked to review (Figure 8). Moreover, the interface can adapt to different question types, by showing different answering fields for each of them. Finally, all information sources that were used to answer the question can also be reviewed. By these means, domain experts can perform an informed assessment.

**Figure 8.**
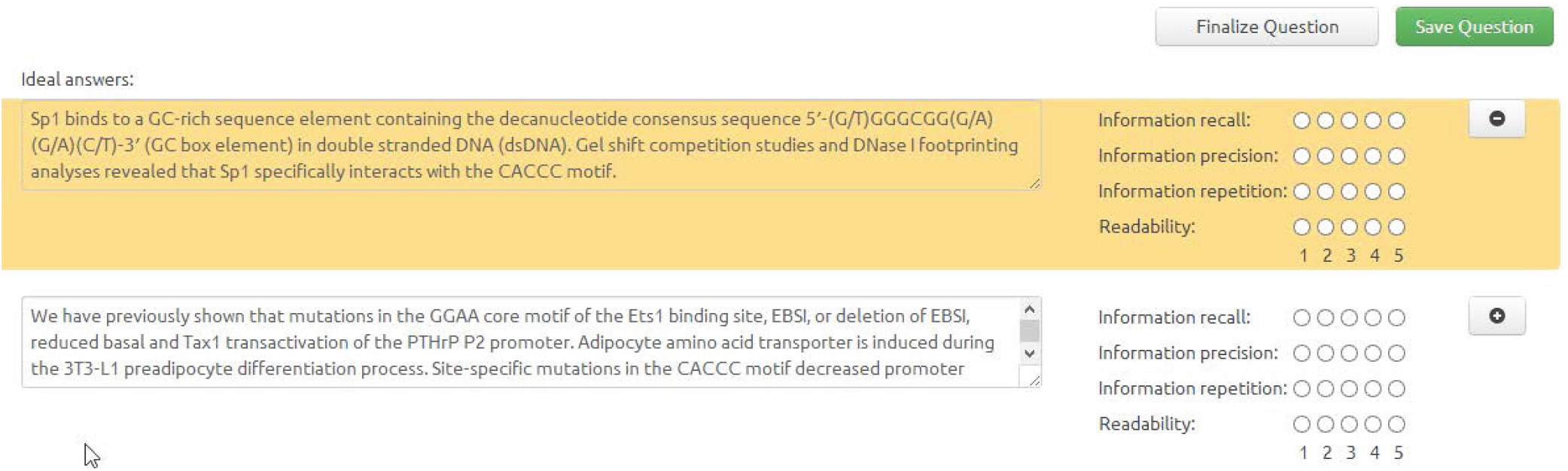
Assessment tool for evaluating system answers. The gold standard answer is at the top.

The assessment tool plays a key role in the creation of the benchmark and the quality assurance of the results generated by the experts during the BioASQ challenges. Moreover, the assessment tool allows the experts to improve their own gold answers and associated material, based on the answers provided by the systems. In particular, the experts revise the documents and snippets returned by the systems, and enrich the gold answers with material identified by the systems. This leads to an improvement of the benchmark datasets that are provided publicly.

## Code availability

BioASQ has created a lively ecosystem, supported by tools and systems that facilitate the creation of the benchmarks. All software is provided with open-source licenses^9^. In addition, the data produced are open to the public^10^.

## Data Records

The dataset follows the *JSON* format. Specifically, it contains an array of questions, where each question (represented as an object in the *JSON* format) is constructed as shown in Table 5.

**Table 5.**
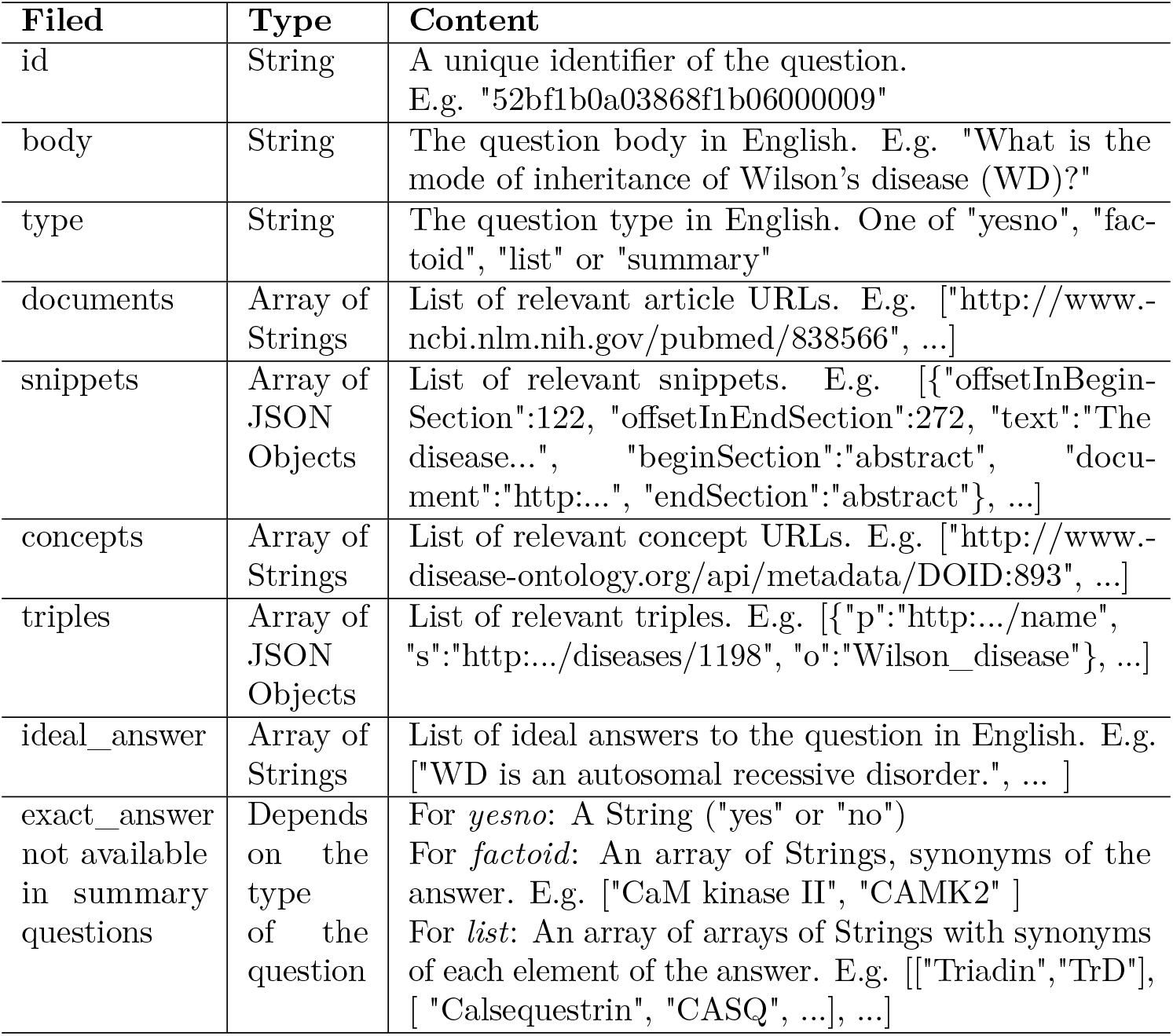
JSON format of the BioASQ-QA benchmark dataset.

## Technical Validation

### Improving the state-of-the-art performance

The participation of strong research teams in the BioASQ challenge has helped to measure objectively the state-of-the-art performance in biomedical question answering. During BioASQ this performance has improved (Figures 10 and 11). It is particularly encouraging that the BioASQ biomedical expert team have assessed the ideal answers provided by the participating systems as being of very good quality. The average manual scores are above 80% (above 4 out of 5 in Fig.11^11^). Still, there is much room for improvement and the future challenges of BioASQ, as well as the benchmark datasets that it provides, will hopefully push further towards that direction.

**Figure 9.**
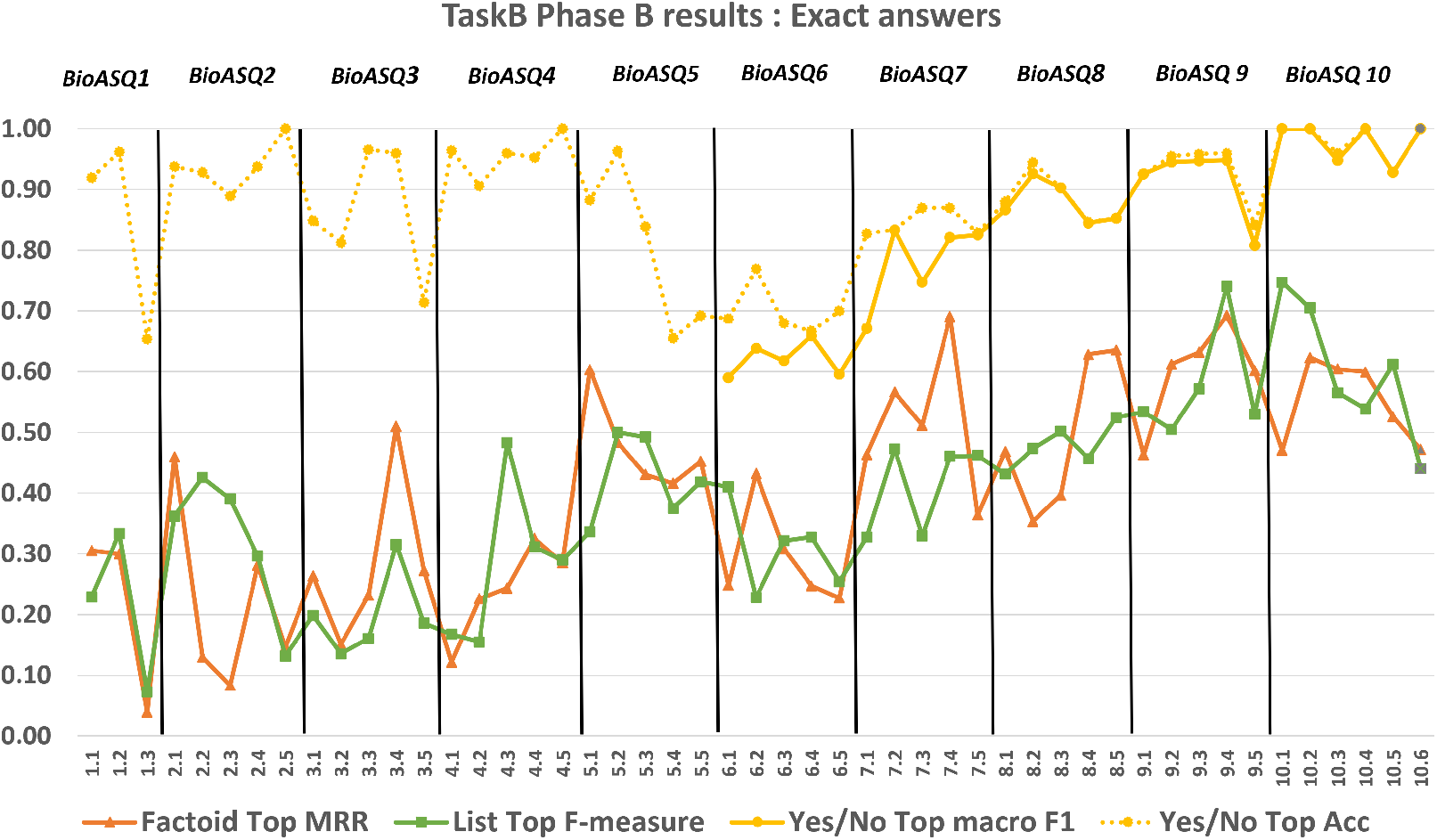
The performance achieved by systems in exact answer generation, across different years of the BioASQ challenge. For each test set the performance of the best performing system (Top) is presented based on the official evaluation measures. Since BioASQ6 the macro-averaged F1 score (macro F1) is the official measure for Yes/No questions, but accuracy (Acc), the former official measure, is also presented.

**Figure 10.**
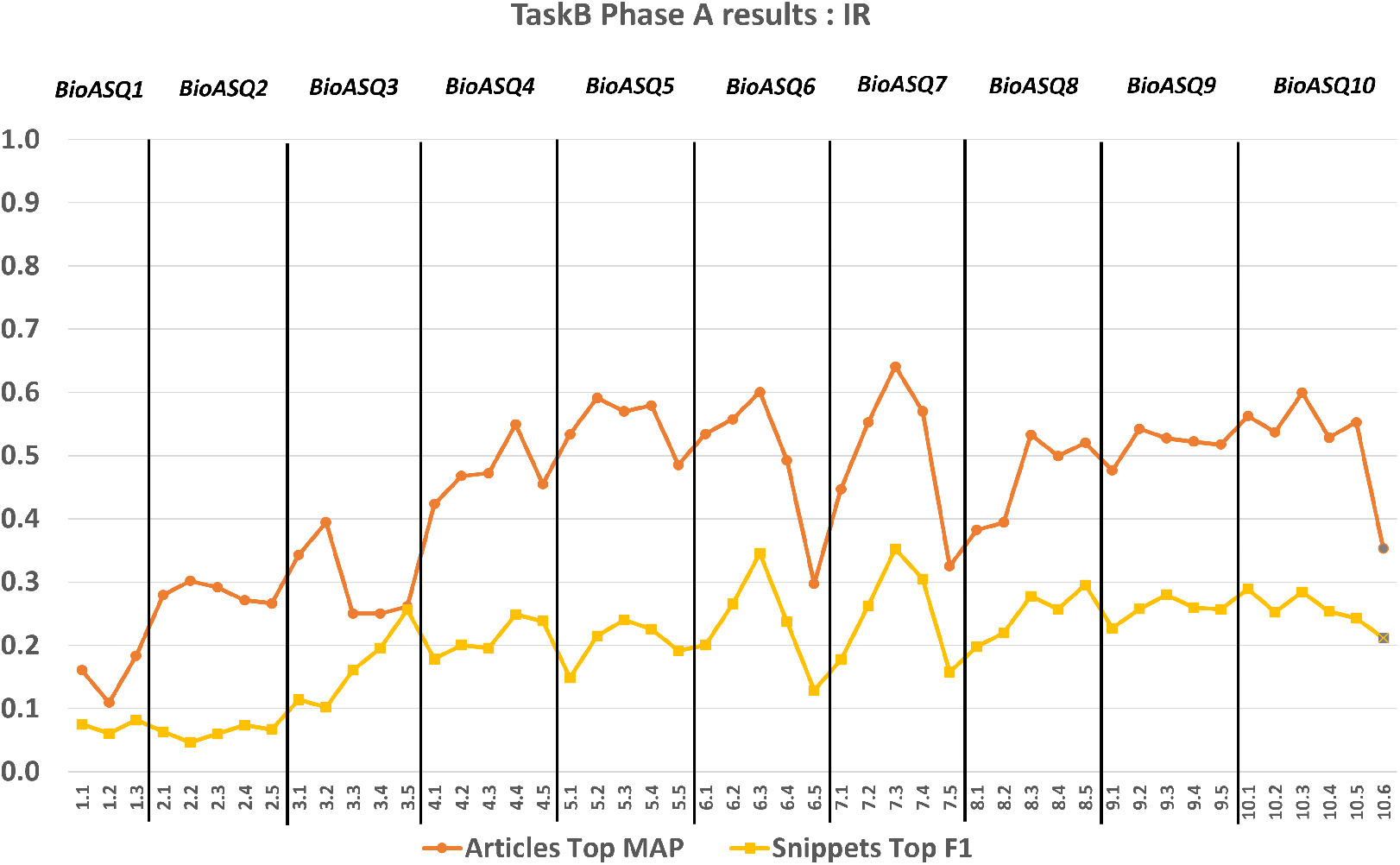
The performance achieved by systems in the information retrieval part of Task B (Phase A), across different years of the BioASQ challenge. For each test set the performance of the best performing system (Top) is presented, based on the official evaluation measures.

**Figure 11.**
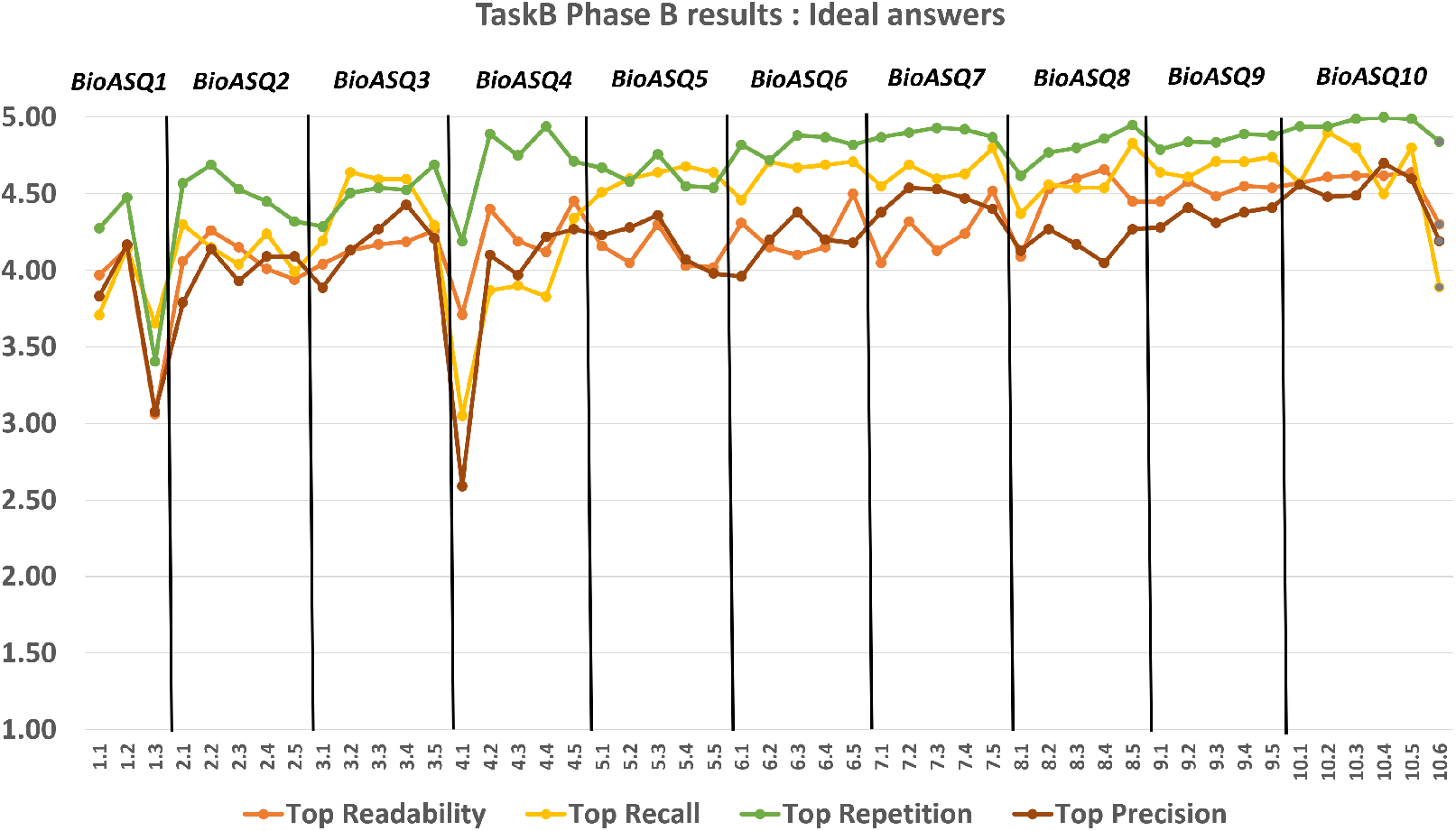
The performance achieved by systems in the ideal answer generation part of Task B, Phase B, across different years of the BioASQ challenge. For each test set the performance of the best performing system (Top) is presented based on the official evaluation measures.

### Usage Notes

Up to date guidelines and usage examples pertaining the dataset can be found in: http://participants-area.bioasq.org/

## Conclusions

This paper presents the BioASQ-QA dataset, providing details about its creations process and the BioASQ ecosystem. The dataset, which is updated yearly, contains currently 4,721 questions and answers, reflecting real information needs of biomedical experts.

Based on the evaluation of the participating systems from the experts, one very interesting result is that humans seem satisfied by “imperfect” system responses. In other words, they are satisfied if the systems provide the information and answer needed, even if it is not perfectly formed.

Another important point is that the BioASQ challenge, as well as the environment in which it takes place, evolve. One consequence of this is the changes that we had to make in the data generation process in response to feedback from the experts and the participants. Additionally, the evolution of vocabularies and databases cause complications. For example, each year’s data are annotated with the current version of the MeSH hierarchy, which is updated annually. In addition, only the articles of the current year of annotation are used for formulating the answers, while articles that will appear in the future may also be of relevance. These are issues that we need to handle and adopt it, in order to have real-life, useful challenge and relevant dataset.

The BioASQ challenge will continue to run in the coming years, and the dataset will be further enriched with new interesting questions and answers.

## Acknowledgements

Google was a proud sponsor of the BioASQ Challenge in 2020, 2021, and 2022. BioASQ was also sponsored by Atypon Systems inc. and VISEO. BioASQ is grateful to the biomedical experts, who have created and manually curated the dataset, as well as to the participants during all these years. Also, BioASQ is grateful to NIH/NLM, who has supported the project by two conference grants (grant n.5R13LM012214-02 and 5R13LM012214-03). For the first two years, BioASQ has received funding from the European Commission’s Seventh Framework Programme (FP7/2007-2013, ICT-2011.4.4(d), Intelligent Information Management, Targeted Competition Framework) under grant agreement n. 318652.Last but not least, BioAQ is grateful to all past collaborators, namely the University of Houston (US), Transinsight GmbH (DE), Universite Joseph Fourier (FR), University Leipzig (DE), Universite Pierre et Marie Curie Paris 6 (FR), Athens University of Economics and Business – Research Centre (GR).

## Author contributions

GP and AK originated the BioASQ challenge and dataset creation. All authors participated in the collection of the data, the development of the annotation and the evaluation tools, and validated the data. AK and AN drafted the manuscript. All authors reviewed the manuscript.

## Footnotes

1 https://www.nlm.nih.gov/bsd/pmresources.html

2 www.BioASQ.org

3 https://www.nlm.nih.gov/mesh/meshhome.html

4 http://linkedlifedata.com/

5 https://www.uniprot.org/

6 http://at.BioASQ.org/

7 See http://www.ncbi.nlm.nih.gov/books/NBK3827/#pubmedhelp.Search_Field_Descrip for a detailed description of the PubMed tags.

8 BioASQ 1 produced only 10 questions, as the first round of the challenge acted as a proof-of-concept. These 10 questions have been incorporated into the BioASQ 2 set.

9 https://github.com/BioASQ

10 http://participants-area.BioASQ.org/

11 In BioASQ 4, the performance was lower due to the fact that it was the first time BioASQ run after the end of the EU-funded project, and the top teams in ideal answer generation started participating after the second batch of the same year.

